# Structural basis of ubiquitin-independent PP1 complex disassembly by p97

**DOI:** 10.1101/2022.06.24.497491

**Authors:** Johannes van den Boom, Hemmo Meyer, Helen Saibil

## Abstract

The AAA+ ATPase p97 (also called VCP, or Cdc48 in yeast) unfolds proteins and disassembles protein complexes in a myriad of cellular processes, but how a substrate complex needs to be loaded onto p97 by a dedicated substrate adapter and then disassembled by p97 has not been structurally visualized so far. Here we present cryo-EM structures of p97 in the process of disassembling a protein phosphatase-1 (PP1) complex by stripping off an inhibitory subunit. We show that PP1 and its partners SDS22 and inhibitor-3 (I3) bind to a peripheral N-domain of p97 via a direct contact between SDS22 and a groove in the N-domain. A density consistent with the SHP box of the p37 adapter binds to the same N-domain underneath the PP1 complex, while the p37-UBX domain is found on the adjacent N-domain. I3 is likely represented by three densities. One covers the PP1 catalytic site adjacent to SDS22, another is at the PP1 binding site for the RVXF motif in I3 pointing towards the p97 pore, and the third is a peptide threaded through the central channel of the spiral-shaped p97 hexamer. Our data show how p97 arranges a substrate complex between the N-domain and central channel, and then extracts one component by threading it through the channel to disassemble the complex.

## Introduction

The AAA+-type ATPase p97 (also called VCP, or Cdc48 in yeast) is critically involved in a large number of diverse cellular signaling and stress response pathways including ER-associated degradation (ERAD), ribosomal quality control, DNA replication and damage repair, protein phosphatase-1 (PP1) biogenesis, and autophagy that together ensure cell survival, proliferation and homeostasis. In all these processes, p97 is believed to unfold substrate proteins in order to dissociate clients from cellular structures or to disassemble protein complexes (Stach and Freemont, 2017; van den Boom and Meyer, 2018; Ye et al., 2017). p97 was shown to be a promising drug target in certain cancers and clinical trials are under way (Anderson et al., 2015; Roux et al., 2021). On the other hand, missense mutations in p97 cause a dominantly inherited multisystem proteinopathy-1 (MSP-1) featuring inclusion body myopathy, Paget’s disease of bone, amyotrophic lateral sclerosis, frontotemporal dementia, and Parkinsonism (Kimonis et al., 2008; Meyer and Weihl, 2014).

Each subunit of the p97 hexamer contains two AAA+ ATPase domains, D1 and D2, that form two stacked hexameric rings enclosing a central channel. The regulatory N-domains are positioned at the periphery of the D1 ring and undergo dynamic up-and-down movements controlled by adapter binding and ATP hydrolysis in D1. In contrast, D2 generates the main driving force for protein unfolding and is essential for cell survival (van den Boom and Meyer, 2018; Ye et al., 2017). Biochemical reconstitution revealed that, as in other AAA+ unfoldases, substrate proteins are inserted in the D1 pore, threaded through the central channel of p97 by the sequential activity of the hexamer subunits and ejected from the D2 pore (Bodnar and Rapoport, 2017; Weith et al., 2018). Cryo-electron microscopy (cryo-EM) analysis showed that the active p97 hexamer is in a staircase configuration with the substrate peptide being transported by sequential interaction with the spiral of pore loops (Cooney et al., 2019; Pan et al., 2021; Twomey et al., 2019; Xu et al., 2022). The pore loops provide a non-sequence-specific interaction between aromatic residues and the peptide backbone that allows it to be threaded through the channel in a hand-over-hand mechanism. ATP hydrolysis in the subunit at the bottom of the spiral results in a discontinuity, or seam, as it triggers the detachment of this subunit from the spiral and induces its reattachment to the top of the substrate peptide. Thus, the peptide is pulled through the pore with the progression of nucleotide binding and hydrolysis around the ring.

Substrate recruitment and engagement are mediated by at least two alternative types of substrate adapters that bind to the N-domains of p97 (Buchberger et al., 2015; van den Boom and Meyer, 2018). A large fraction of p97 substrates are ubiquitylated. The Ufd1-Npl4 adaptor binds the ubiquitin chain and inserts one ubiquitin into the D1 pore leading to threading of the attached substrate through the channel for substrate unfolding (Twomey et al., 2019). In contrast, ubiquitin-independent substrate targeting is mediated by adapters of the SEP-domain family such as p37 (also called UBXN2B). Most prominently, p97-p37 regulates the biogenesis of PP1 holoenzymes by disassembling a regulatory complex of PP1 catalytic subunit (called PP1 from here on) with its partners SDS22 (also called PPP1R7) and Inhibitor-3 (I3, also called PPP1R11) (Weith et al., 2018). Disassembly is achieved by threading I3 through the p97 channel which strips I3 (and SDS22) off PP1 to allow PP1 holoenzyme formation with other PP1 subunits (Weith et al., 2018). Notably, I3 targeting relies on multivalent recognition of the SDS22-PP1-I3 (SPI) complex by the p37 adapter (Kracht et al., 2020), and involves an internal recognition site within I3 that is inserted as a loop into the p97 pore to initiate unfolding of I3 (van den Boom et al., 2021).

While the substrate threading process has been well characterized, it is unclear how substrate proteins are recruited and positioned for disassembly so that the substrate can be delivered into the pore. This is particularly important in protein complex disassembly reactions when one subunit is inserted into the p97 pore while other subunits are spared. Moreover, no structural information exists on how ubiquitin-independent targeting to p97 by the p37 adaptor is achieved. Therefore, the mechanism of PP1 complex disassembly and protein complex disassembly in general is not understood.

Here we have analyzed assemblies of p97 with its p37 adapter and the SPI substrate complex in the presence of ADP-BeF_x_ by cryo-EM, yielding a set of structures of p97-p37 during the disassembly of the SPI substrate. Unexpected findings provide new insights into the spatial arrangement of a ubiquitin-independent substrate and the adapter protein on p97 that facilitates complex disassembly.

## Results

### Human p97 in the act of translocating a specific substrate protein

To obtain structural data on how p97-p37 disassembles the SPI complex, we added an excess of p97 and p37 to the SPI complex. Substrate engagement was induced by incubation in the presence of ADP and beryllium fluoride (ADP-BeF_x_). The resulting p97-p37-SPI complex was purified by size exclusion chromatography and analyzed by cryo-EM. Our structures show p97 loaded with the SPI complex at a defined position on the N-domain in the act of translocating a segment of I3. The density map (Figure 1a-c and Supplementary Figure S1a) confirms a conserved peptide threading mechanism previously proposed for p97 and yeast Cdc48 (Cooney et al., 2019; Pan et al., 2021; Twomey et al., 2019; Xu et al., 2022). Both D1 and D2 domains of p97 are in the staircase configuration with subunit A on top, followed by subunits B, C and D in a clockwise manner to subunit E at the bottom (Figure 1a and b). Subunit F bridges the bottom subunit E with the top subunit A and has weaker electron density (Supplementary Figure S1b), likely reflecting its flexible position. All six N-domains of p97 are in the up-conformation, albeit the resolution for the A and F subunits is lower than for subunits B-E (Supplementary Figure S2). The central pore is filled in both ATPase rings with a density of an extended polypeptide peptide strand (Figure 1b) that is engaged for threading via interactions to the D1 and D2 domains of five p97 subunits (A to E).

**Figure 1.**
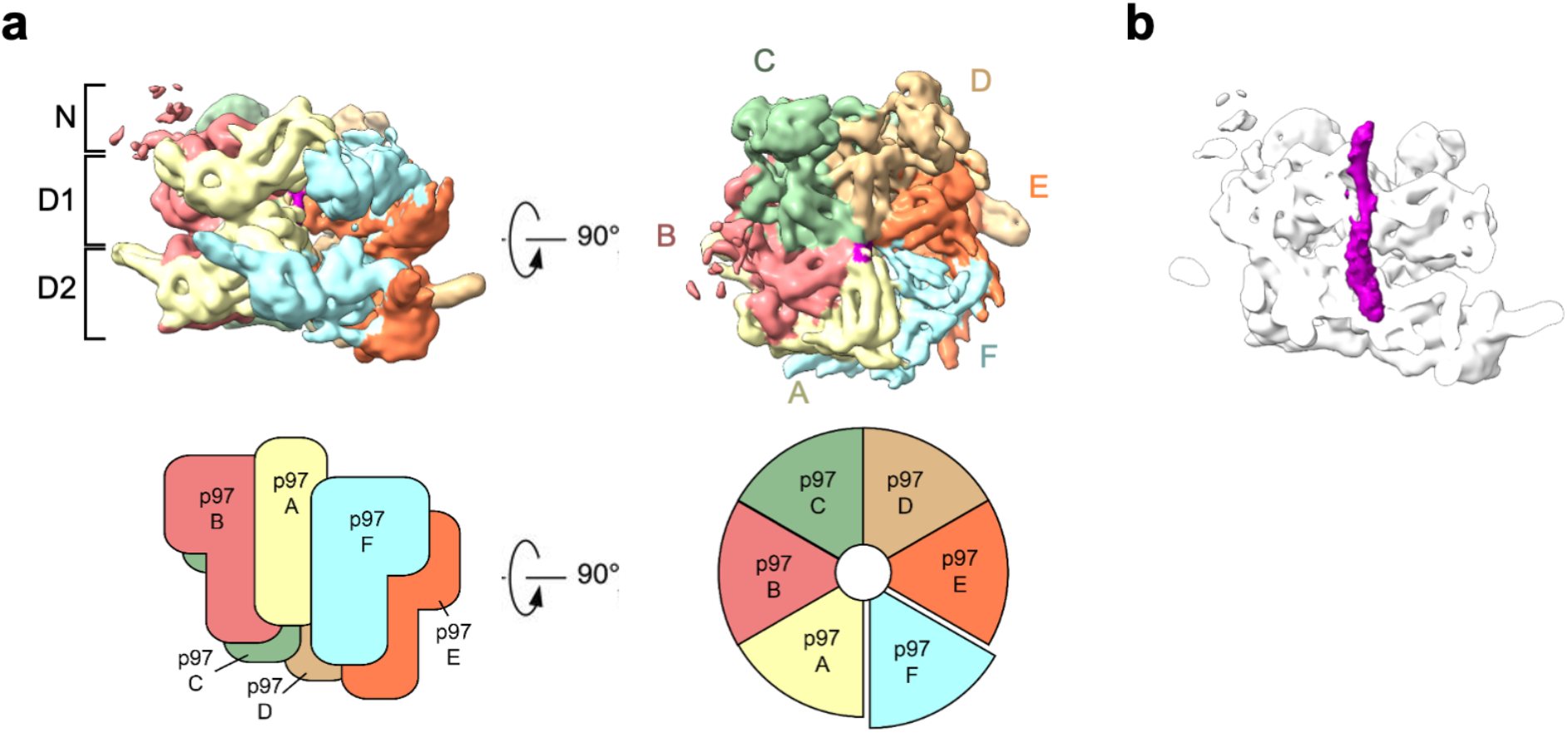
Structure of human p97 during substrate translocation. (a) Side (left) and top (right) views of the of the cryo-EM map of the p97 hexamer (0.00851 threshold). In the top view, subunit A, pale yellow, is followed by subunits B to F arranged in a clockwise staircase. Schematic views of the subunit assembly are shown in the lower panel. (b) Side view section of p97 showing the substrate density (purple) in the central channel.

There is previous biochemical evidence that p97 initiates the translocation of I3 from a position close to the N-terminus of the protein, taking a short peptide hairpin of I3 through the pore (van den Boom et al., 2021). Our map shows density corresponding to a single strand of I3 within the D1 ring pore, while the density inside the D2 ring pore is wider and compatible with a hairpin loop of I3 (Figure 1b). However, the resolution of the density for the mobile and disordered substrate is insufficient to reliably distinguish between single and double peptide strands.

### The SPI substrate complex is recruited to the N-terminal domain of p97

Our image dataset was sorted into classes yielding maps representing different populations of SPI-loaded p97 complexes, with SPI bound to N-terminal domains of p97 subunits at different positions in the hexameric staircase. We obtained four individual maps, in which the SPI complex is positioned on the second, third and fourth highest p97 subunit (B, C, D and E subunit), respectively (Figure 2a and b and Supplementary Figure S3). The SPI densities occupy a consistent position above the different p97 N-terminal domains with weaker densities extending slightly towards the channel entrance near the central axis of p97. This suggests that the SPI substrate complex remains bound to the same p97 subunit which then, together with SPI, proceeds through the different positions in the staircase configuration (A-F) during the translocation cycle. Subunit B shows the best-defined SPI density, while the density becomes more disordered as it advances through the cycle until subunit E (Figure 2a and b). We therefore focused our analysis on the structure with SPI bound on the B subunit of p97.

**Figure 2.**
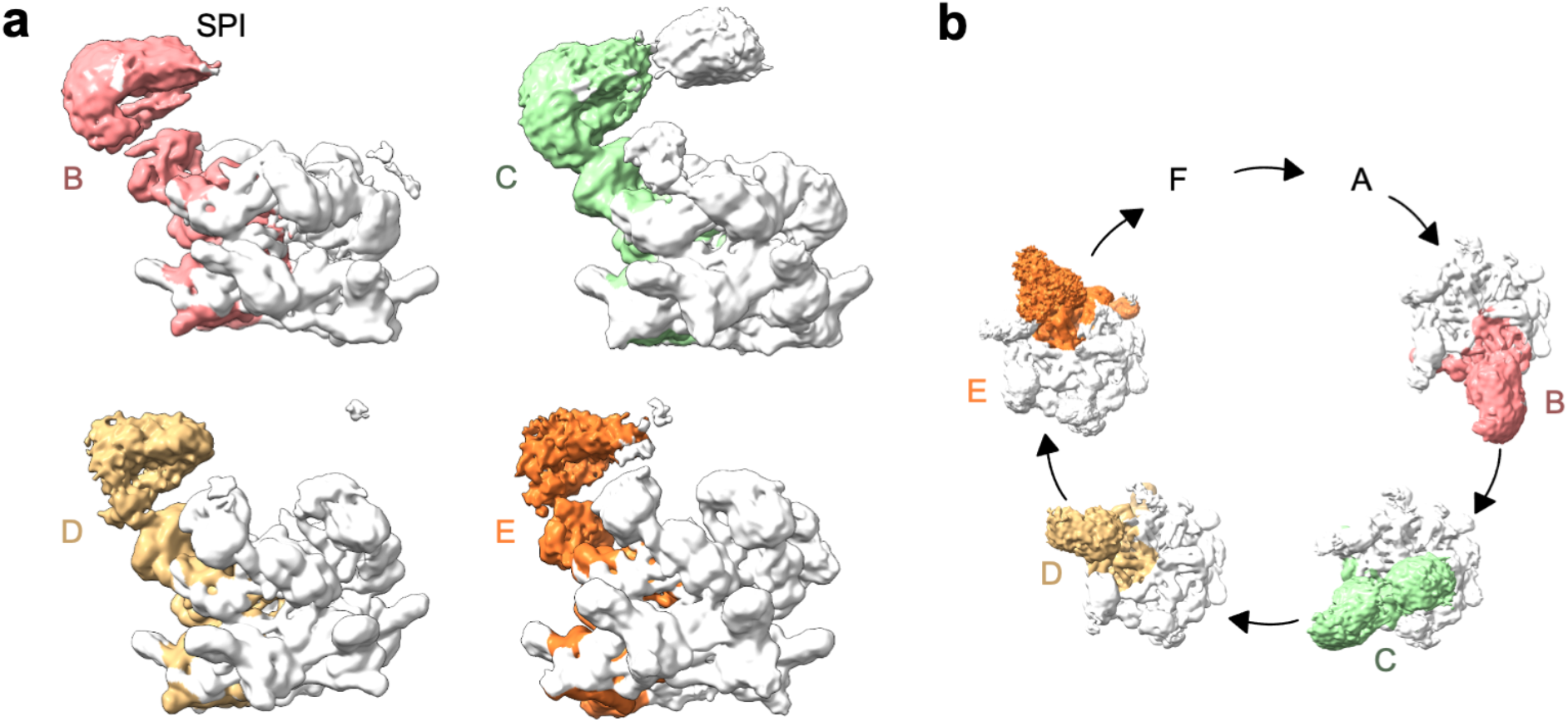
p97-p37 loads the SDS22-PP1-I3 (SPI) substrate complex onto its N-terminal domain. (a) Cryo-EM maps of four different structural classes show density of one SPI complex above the N-terminal domain in p97 subunits B, C, D or E, respectively. An extra density (white) above the channel was seen for the subunit C complex, which did not include enough images to group this feature into a separate class, as was done for the other subunit complexes. (b) Top views of SPI complex density bound to subunits B-E. Subunits F and A showed weak density for SPI due to the flexibility of their interface. Density thresholds: B, 0.00749; C, 0.00363; D, 0.0032; E, 0.00241

### SDS22 docks in the p97 N-domain groove while p37 spans two adjacent N-domains

The SPI complex on subunit B is seen very clearly as an arc-shaped density enclosing a globular density (Figure 3a-c), corresponding well with the available X-ray crystal structures of the PP1-SDS22 complex, in which the leucine-rich repeats (LRRs) of SDS22 form an arc around PP1 (Choy et al., 2019). It is generally assumed that substrates are recruited to the p97 N-domain indirectly through substrate adapters. Surprisingly, our structure shows a direct interaction of SDS22 with the N-domain of p97 (Figure 3b). Closer analysis of this interface revealed docking of a short helix (aa 337-346), located C-terminally of the LRR array in SDS22, to the groove between the N_N_ and N_C_ subdomains of the p97 N-domain (Figure 3b, zoomed-in view). We validated this interaction with point mutations in the p97 N-domain groove (G54K or Y143A) that substantially reduced SPI but not p37 binding to p97 (Supplementary Figure S4).

**Figure 3.**
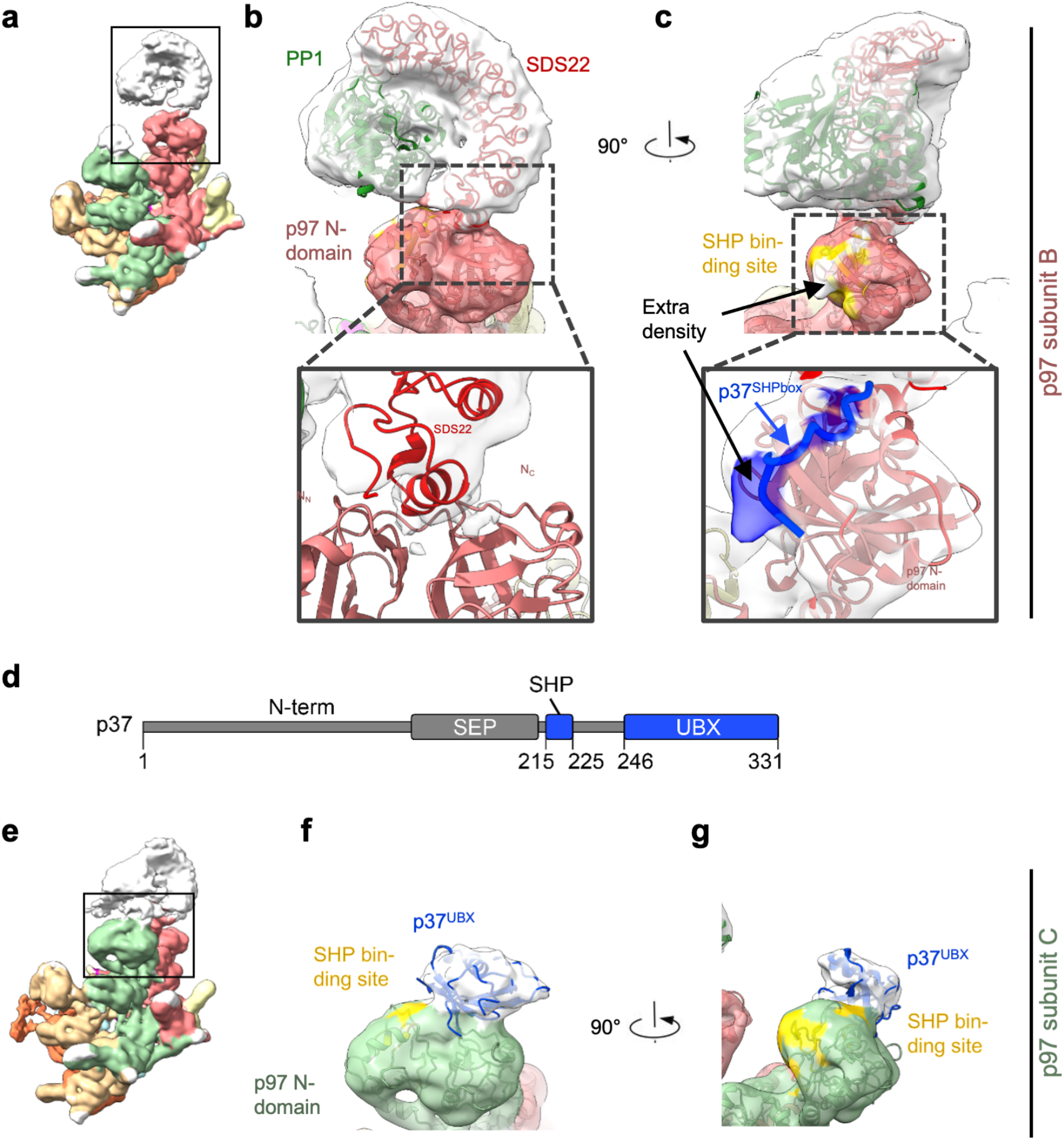
Position of SDS22 and p37 on the p97 N-domains. (a) Shows the complex with the SPI substrate loaded on the B subunit of p97. (b) SDS22 interacts with the N-domain of p97 (subunit B) via its C-terminal helix and binds PP1 on its entire concave face. Zoom-in view shows the interface between SDS22 and the p97 N-domain. SDS22 binds with its C-terminal helix in the groove between the N_N_ and N_C_ lobes of the p97 N-domain. (c) Rotated view showing the extra density (white) at the position of the SHP box binding site (yellow). Zoom-in view shows the modelled SHP box peptide (blue). (d) Domain composition of the p37 adapter. N-term: unstructured N-terminus; SEP: SEP (Shp1, eyes closed, p47) domain; SHP: SHP box for p97 binding; UBX: ubiquitin regulatory X (UBX) domain for p97 binding. Parts resolved in the structure are depicted in blue. Residue numbers of structural elements are indicated. (e) Shows the same map as in (a) viewed with the C subunit of p97 in front. (f) The N-domain of the p97 C subunit (green) adjacent to the SPI-bound B subunit binds the UBX domain of p37 (blue). (g) The binding site for the p37 SHP box (yellow) is not occupied on the p97 C-subunit. Density threshold of the map is 0.0024.

Substrate adaptors such as p37 and its paralogues, as well as Ufd1-Npl4, interact through two different anchor points with the p97 N-domain, a linear SHP box and a UBX domain (Figure 3d) (Bruderer et al., 2004). While previous structures show these interactions individually (Cooney et al., 2019; Dreveny et al., 2004; Hanzelmann and Schindelin, 2016; Le et al., 2016; Li et al., 2017; Lim et al., 2016; Xu et al., 2022), it has remained controversial whether the SHP box and UBX domain dock to the same or different N-domains within the p97 hexamer. In our structure, there is extra density on the p97 N-domain at the SHP box binding site underneath the PP1 complex, which well accommodates the SHP box peptide from the p97-Ufd1 structure (Figure 3c) (Le et al., 2016). Of note, the UBX domain of the adapter cannot bind the same N-domain as its interaction site in the N-domain groove is occupied by SDS22. Instead, we see a UBX domain on the adjacent subunit (Figure 3e-g) with its N-terminus facing towards the SHP box peptide. The distance between the SHP box peptide and UBX domain on the adjacent N-domains is 50 Å, which can easily be bridged by the 20 residues in the linker between SHP box and UBX domain. Therefore, our structural results point to an extended structure for the p37 adaptor, in which the SHP box peptide and UBX domain bind to successive p97 subunits. This configuration locates the linker underneath PP1, which is consistent with our previous finding that the linker is critical for interaction with PP1 and recruitment of the SPI complex (Kracht et al., 2020).

### Assignment of additional I3 densities on PP1

The SPI density above the N-domain of p97 contains two regions of additional density that are not filled by PP1 or SDS22 (Figure 4) suggesting they are occupied by I3. This is consistent with two binding sites of I3 to PP1 (Zhang et al., 2008). One site of unassigned density is adjacent to the PP1 active site (Figure 4 density 1). Binding of this region by a C-terminal part of I3 (aa 65-77) is critical for the inhibitory activity of I3 for PP1 (Zhang et al., 2008). The other is on the opposite side of PP1, adjacent to the binding site in PP1 for the RVXF motif of I3 (aa 40-43 in I3), which is essential for I3-PP1 interaction (Figure 4 density 2). The distance between these two sites is approximately 30 Å which is compatible with the 21 residues between the two regions in I3 (Zhang et al., 2008). This suggests that the RVXF motif and the inhibitory region of I3 account for the extra densities, while other parts of I3 which are not interacting with PP1 are likely to be disordered and therefore not visible in the density. The distance between the density at the tip of the SPI complex corresponding to the RVXF motif in I3 and the p97 pore that accommodates the proposed inserted N-terminal section of I3 is approximately 60 Å (Figure 5a and b). This is compatible with the roughly 40 amino acids N-terminal of the RVXF motif that are first targeted for pore insertion (van den Boom et al., 2021).

**Figure 4.**
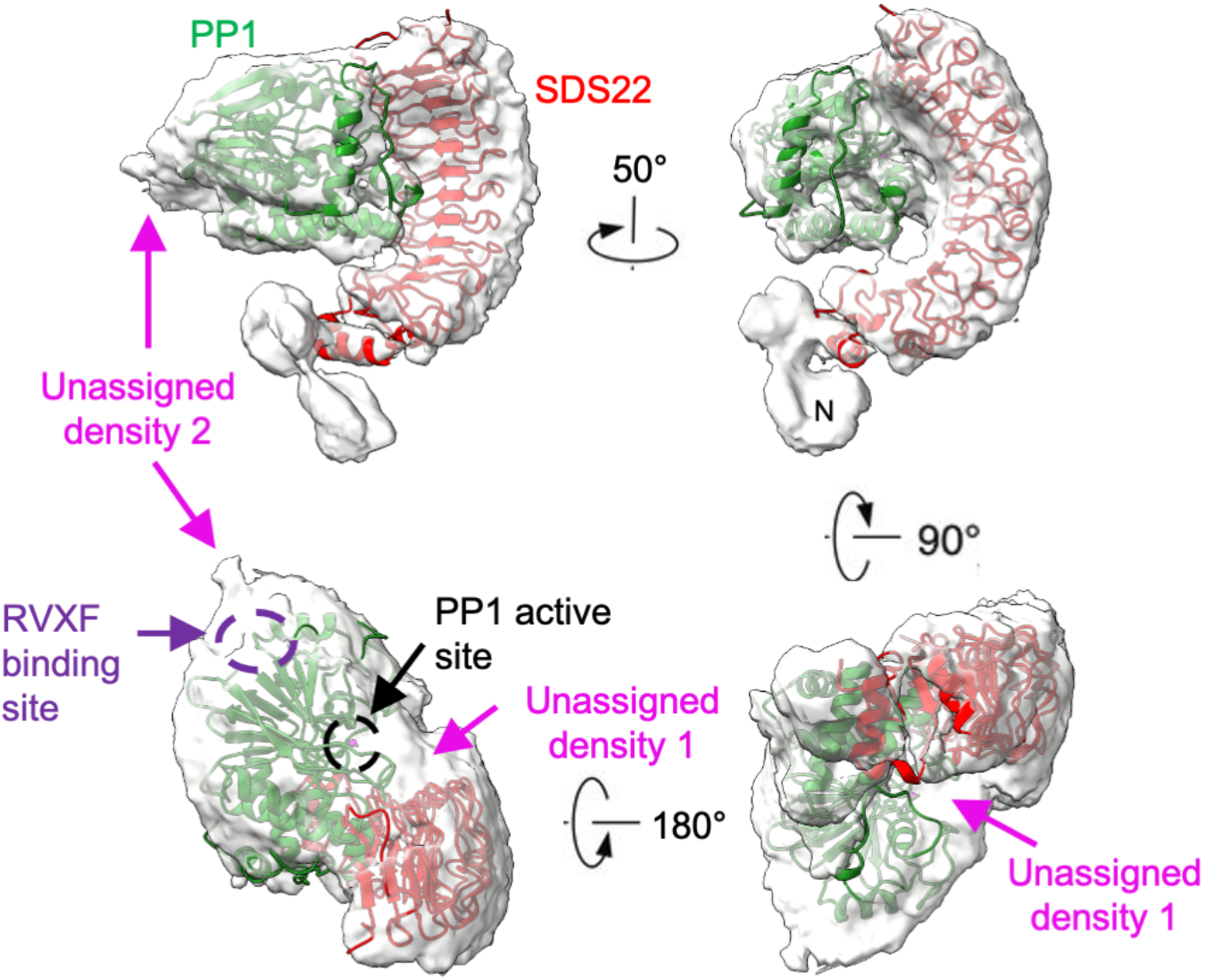
Structure of the SPI complex with p97. Side, front, top and bottom views the SPI complex of subunit B (threshold 0.00354) with fitted crystal structures of PP1 and SDS22 (PDB: 6obn). Additional density at the bottom of the upper panels is part of the N-domain. There are two regions of unassigned density between PP1 and SDS22. Density 1 is next to the active site of PP1 (black dashed circle). Density 2 is adjacent to the binding site for the RVXF motif in I3 (purple dashed circle). The unassigned densities likely represent parts of I3.

**Figure 5.**
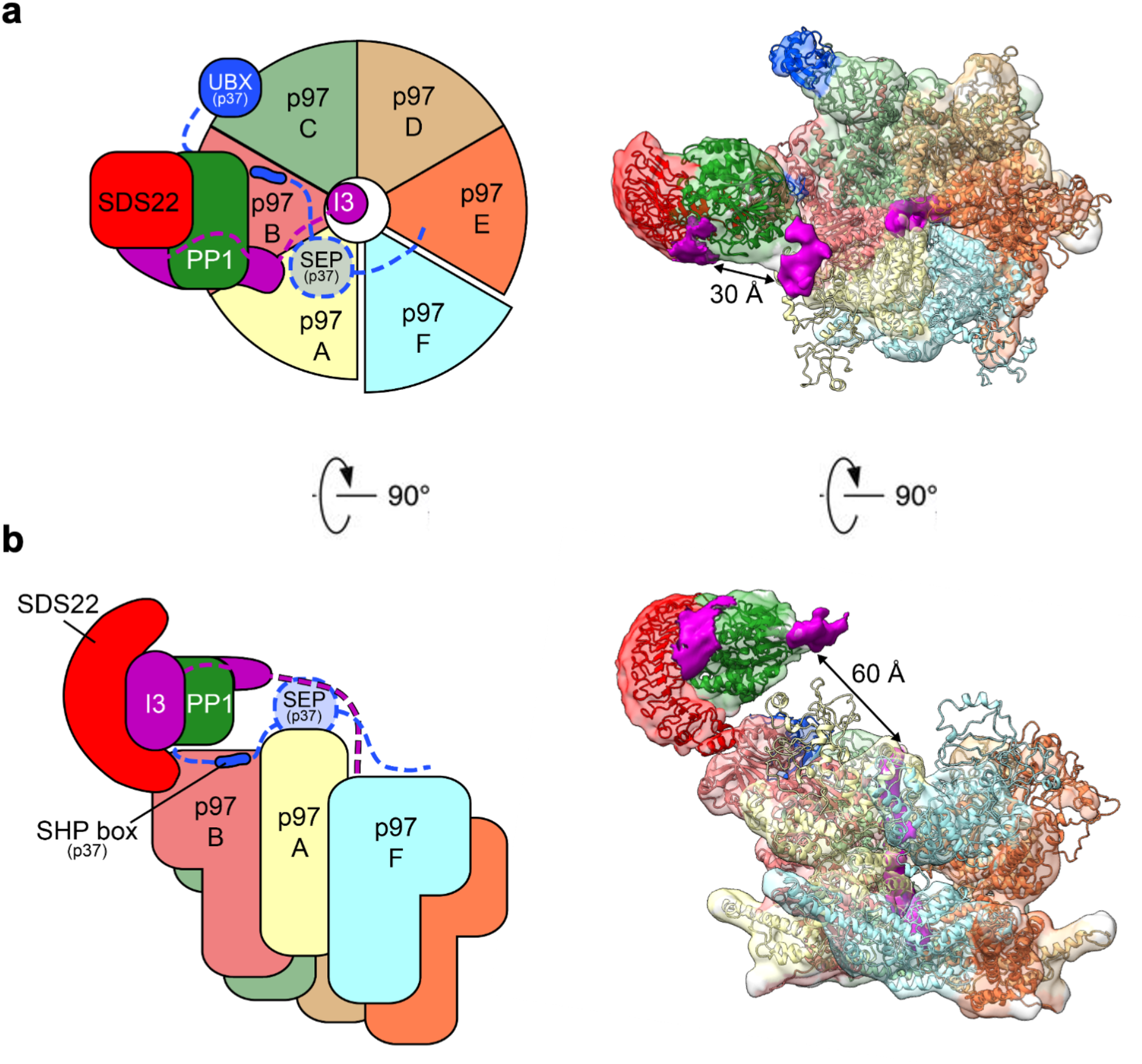
Schematic of proposed p97 processing of the SPI complex. (a) Top view of p97-p37 during disassembly of the SPI complex as cartoon (left panel) and density map with models (right panel, threshold 0.0034) in top (top panel) and side view (bottom panel). The p97 hexamer (subunits A-F) is in staircase conformation with subunit A on top and E on the bottom. The p37 adapter positions the SPI on the p97 B subunit of the p97 hexamer (subunits A-F) with the p37 SHP box bound to the B subunit and the p37 UBX domain bound to the p97 N-domain of the adjacent C subunit. The SPI complex is bound on the p97 hexamer via a direct interaction of the SDS22 C-terminal helix with the p97 N-domain of the B subunit. The concave side of SDS22 faces towards the p97 pore and is occupied by PP1. The unfolded polypeptide in the central channel of the p97 hexamer is likely the N-terminal part of Inhibitor-3 (I3) and interacts with the p97 subunits A-E, while the F subunit is disengaged. The RVXF motif of I3 is attributed to the electron density at the RVXF binding site of PP1, located 60 Å above the p97 pore and possibly guided to the pore by the p37 SEP domain. The electron density covering the active site of PP1 likely reflects the inhibitory region in the C-terminal part of I3.

## Discussion

Using the transition state intermediate ADP-BeF_x_ to trap the p97 complex during its action on the SPI substrate, we have determined multiple structures of p97 with the SPI substrate bound to subunits in different stages of the ATPase cycle (Figure 2). The SPI structure is best resolved on subunit B, making that state the most informative. Previous structural studies have used cryo-EM to reveal the spiral conformation of p97 with a substrate trapped in the pore (Cooney et al., 2019; Pan et al., 2021; Twomey et al., 2019; Xu et al., 2022), but have not revealed a loading complex with the adaptor proteins delivering the substrate to p97 for protein complex disassembly. Our structure of p97 with a loaded substrate complex on the N-domain reveals unanticipated new principles of how a substrate is positioned and engaged onto p97 with the help of the adapter leading to dissociation of one subunit while sparing the other components.

Perhaps the most prominent new feature of our structure is the precise docking of the SPI substrate complex on the p97 N-domain. This is suggested by the well-defined structure of SDS22-PP1 and its interaction interface with the N-domain. Surprisingly, SDS22 appears to make direct contact with the groove between the N_N_ and N_C_ lobes of the p97 N-domain that can otherwise accommodate a loop of the UBX domain of the adapter. This is unexpected because it has been assumed that substrate specificity is conferred solely by the adapter. This binding mode excludes binding of the UBX domain of the adapter to the same N-domain. In fact, while the structure shows a density corresponding to the p37 SHP box underneath the SPI complex, it also reveals a density well corresponding to the UBX domain on the adjacent N-domain. The distance can be readily bridged by the linker between SHP box and UBX domain, which however is not visible in our structure. The proposed position of the bridging linker is also consistent with previous data on a crosslink between the linker and PP1 (Kracht et al., 2020). Thus, the specific docking of the SPI onto the N-domain appears to be a result of a multivalent contribution of adapter, the p97 N-domain and subunits of the SPI complex.

The unassigned densities not filled by SDS22 or PP1 on the SPI complex correspond well with biochemically established I3 binding sites covering the catalytic site of PP1 and the RVXF interaction site in PP1 (Zhang et al., 2008), although we cannot exclude that the disordered N-terminus of SDS22 also contributes to this density. Of note, the RVXF site points towards the D1 pore of p97 that harbors an inserted peptide apparently derived from I3. We previously demonstrated biochemically that a sequence element in I3 N-terminal to the RVXF motif is recognized and engaged in the pore, and that this can occur as a short hairpin loop (van den Boom et al., 2021). Given that the distance between these densities is compatible with the 40 residue length of that sequence, it is tempting to speculate that these elements are connected, and that our structure represents I3 engaged in the channel and in the process of being pulled off PP1 (Model Figure 5a and b). The ATPase rings in our structure are in a spiral conformation primed to thread the peptide through the channel by the established hand-over-hand mechanism (Cooney et al., 2019; Pan et al., 2021; Twomey et al., 2019; Xu et al., 2022).

The peptide which connects densities of I3 on PP1 and in the channel is likely flexible and therefore not visible in the density. Of note, previous crosslink experiments suggested that the p37 adapter also binds I3 and that this is mediated by its SEP domain (Weith et al., 2018). As the SEP domain is also not visible in the structure, we speculate that the SEP domain mediates the interaction to the flexible part of I3 in the space between PP1 and I3 in the p97 pore, which is in line with extra density in this region seen at lower density threshold (data not shown).

The well-defined position of the substrate on the N-domain is likely important for dissociation of partners for which one needs to be fixed while the other is extracted. Immobilization of a substrate complex may more generally apply to protein complex disassembly. This concept may even extend to disassembly reactions triggered by ubiquitin and mediated by the alternative adapter Ufd1-Npl4 such as the disassembly of the Ku70/80 complex or the MCM2-7 helicase (Mukherjee and Labib, 2019; van den Boom et al., 2016). It would therefore be interesting to establish whether these protein complexes also bind directly to the N-domain. In contrast, a monomeric ubiquitylated substrate protein would not require immobilization for unfolding, which could explain why a direct interaction of the substrate and the p97 N-domain has not been observed in previous cryo-EM analyses.

The p97 N-domains can undergo up and down movements relative to the AAA barrel, driven by the ATP cycle in the D1 domain (Tang et al., 2010). It is noteworthy that in our structure, all N-domains are in the up position which brings the substrate complex close to the central p97 pore. Given that the pulling forces generated by substrate threading in the channel are believed to strip I3 off PP1 and that the SDS22-PP1 complex is tightly anchored on the N-domain, additional movement of the N-domain could contribute to the forces needed to dissociate a protein from its partner. Further structural and functional studies are needed to clarify these questions.

## Acknowledgements

H.S. thanks the Wellcome Trust for research support (106249/Z/14/Z). Data collection at Birkbeck was supported by Wellcome Trust (202679/Z/16/Z and 206166/Z/17/Z). H.M. thanks the DFG for grant Me1626/3-3. We acknowledge Diamond Light Source for access and support of the cryo-EM facilities at the UK’s national Electron Bio-imaging Centre (eBIC) [under proposal EM 20287], funded by the Wellcome Trust, MRC and BBRSC. We are grateful to Jesus Gomez de Segura for EM and image processing work. We thank Natasha Lukoyanova for Krios data collection and David Houldershaw for computing support.

## Methods

### Sample purification

p97 and the complex of SDS22, PP1γ and His-tagged I3 (SPI complex) were generated in insect cells as previously described (Weith et al., 2018). In brief, Sf9 cells were transfected with a pFL vector encoding His-tagged p97, or His-tagged I3, or both untagged SDS22 and PP1γ using the dual expression cassette. After virus amplification in Sf9 cells, final protein expression was performed in High Five cells. For the expression of the SPI complex, High Five cells were transduced with both viruses (SDS22+PP1 and His-I3).

The SPI complex was purified from cleared High Five cell lysates using a HisTrap FF column (Cytiva) in p97 buffer (50 mM HEPES pH 8.0, 150 mM KCl, 5 mM MgCl_2_, 5% glycerol) with 25 mM imidazole. Protein was eluted directly onto a HiTrap Q HP anion exchange column (Cytiva) with p97 buffer supplemented with 300 mM imidazole. SPI was eluted from the Q HP column using p97 buffer with 1 M KCl and further purified by gel filtration on a Superdex 200 16/600 column (Cytiva) in p97 buffer with 1 mM DTT.

His-p97 was purified identically to the SPI complex via NiNTA affinity chromatography and anion exchange chromatography. However, the protein was eluted from the Q HP column using a gradient (buffer A: p97 buffer; buffer B: p97 buffer + 850 mM KCl), before the buffer was exchanged by repeated concentration and dilution in p97 buffer with 1 mM DTT. Human p37 was generated from GST-p37 in *Escherichia coli* BL21 (DE3) as previously described (Weith et al., 2018). In brief, cleared bacterial lysate was loaded onto a GSTrap column (Cytiva) using p97 buffer, incubated with PreScission protease for GST-tag removal and further purified by gel filtration on a Superdex 200 16/600 column in p97 buffer with 1 mM DTT. Proteins were snap frozen in liquid nitrogen and stored at -80 °C.

### GST-pulldown assays

Binding assays were carried out as described in (Weith et al., 2018). Mutations in p97 were introduced by QuikChange mutagenesis. His-tagged p97 and GST-p37 were generated in bacteria. SPI was made in insect cells as described above.

### Cryo-EM sample preparation

The complexes were reconstituted from frozen aliquots, thawed at room temperature. p97 (5 μM), p37 (4 μM) and SPI (1.5 μM) were mixed in buffer containing 50 mM HEPES pH 7.5, 150 mM KCl, 5 mM MgCl_2_, 2% glycerol, 1 mM DTT, 0.0005% NP40 and 2 mM ADP-BeF_x_. ADP-BeF_x_ was prepared freshly from frozen stocks of ADP (0.1 M), BeSO_4_ (1 M), KF (1 M) and MgCl_2_ (1 M) mixed sequentially in that order in relative volumes 1:1:8:1. After mixing, the sample was incubated at room temperature for 10 min and diluted 3:1 in buffer before plunge freezing in liquid ethane using a Vitrobot Mk IV (Thermo-Fisher Scientific). 4 μl of sample was loaded on freshly glow discharged 1.2-1.3 300 mesh C-flat™ grids without further support (Electron Microscopy Sciences). The Vitrobot chamber was set to 4 °C and 95% humidity and grids were prepared with a range of blotting time and force, from which the best grids were selected after manual screening.

### EM data collection

Micrograph collection was done at Birkbeck College London using a Titan Krios G3i electron microscope (FEI/Thermo Fisher) operated at 300 kV equipped with a post-GIF K3 detector and an energy slit 20 eV wide. The images were taken in super resolution mode at a nominal magnification of 81000 with a pixel size of 0.5335 Å, dose rate of 15.94 e-/Å^2^/s over 3 seconds of exposure and a defocus range from -3.3 to -1.8. 6000 movies were collected using these settings. Preliminary data for this project was obtained in the electron Bio-Imaging Center (eBIC) at Diamond Light Source using a Titan Krios G3i electron microscope (FEI/Thermo Fisher) operated at 300 kV equipped with a post-GIF K3 detector. This data was recorded with a 0.845 Å pixel size and 18.5 e-/Å^2^/s over 1.8 seconds.

### Image processing

For processing, data were resampled to 1.1 Å/pixel. Initial image processing was done in Relion (Scheres, 2012). Movies were motion corrected using Motioncorr2 (Zheng et al., 2017) and CTF was corrected using Gctf (Zhang, 2016). After extraction and preliminary visual evaluation 6 million particles were exported to Cryosparc 2 (Punjani et al., 2017) for 2D classification. After two rounds of classification 1.6 million particles were selected for ab initio reconstruction and non-uniform refinement. The particles and resulting map were exported into Relion for further 3D classification and refinement, yielding a subset of 850000 particles showing signs of substrate occupancy which were selected for further processing. This subset was subjected to two cycles of non-uniform refinement and heterogeneous refinement in Cryosparc, resulting in a final selection of 366000 particles that showed clear densities for the cofactors and substrates. This subset was further processed with Relion for extensive classification. Different masks corresponding to the different possible substrate binding zones were used to obtain subsets showing predominant densities in each of those zones, allowing the study of different populations of SPI bound p97. These masks were created using Relion from densities manually extracted from the maps with UCSF Chimera (Pettersen et al., 2004). Final maps were refined in Relion following the gold standard procedures and presented unmasked, aside from the structure focused on p97 alone, without further post-processing.

Figures were made using UCSF Chimera and ChimeraX (Pettersen et al., 2021). The human homology model was created with Swissmodel (Waterhouse et al., 2018) and based on the structure by (Cooney et al., 2019), then refined in flexEM (CCPEM) (Topf et al., 2008). Real space refinement in COOT was used to optimize the fit (Emsley et al., 2010). For focused flexEM refinements and cross-correlation measurements a segment of the map corresponding to the SPI and UBX densities was extracted using the zone function in UCSF Chimera. SPI subunit models were based on the structure by (Cooney et al., 2019).

