## Supplementary figures for "Structural basis of ubiquitin-independent PP1 complex disassembly by p97"

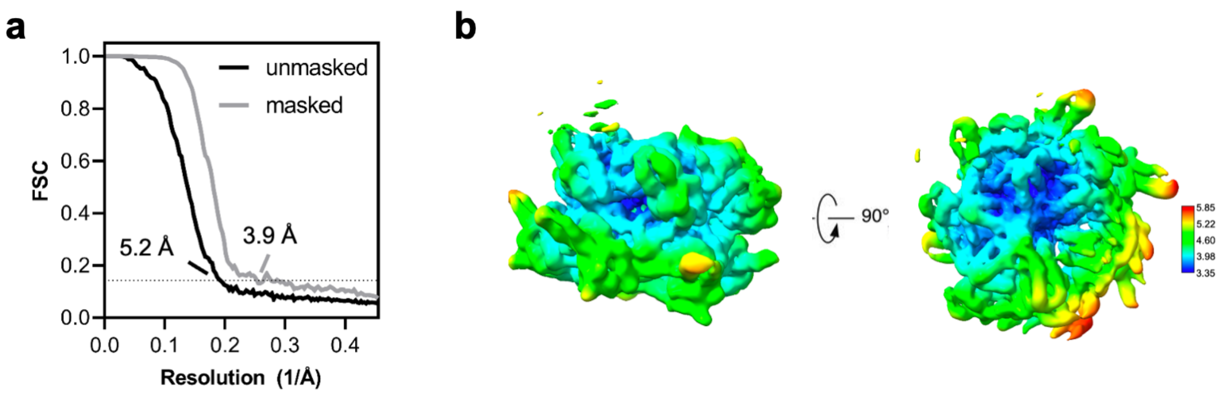


**Figure S1.**

(a) Gold-standard Fourier shell correlation (GSFSC) calculated during refinement without and with mask. The resolutions were determined at FSC = 0.143 (dotted line). (b) Local resolution was calculated from the half-map and colored according to the scale on the side. For this map 866937 particles were used.

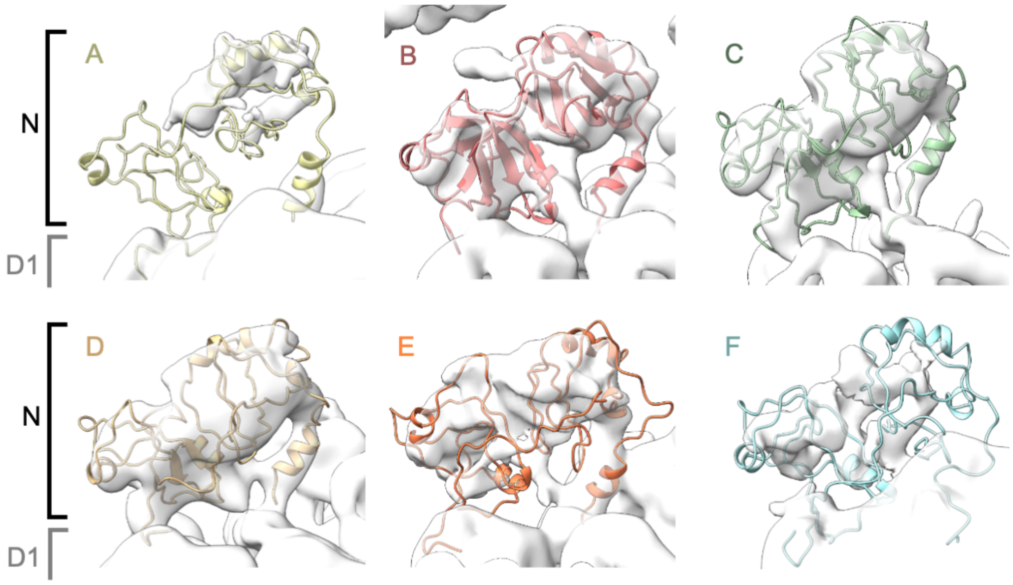


**Figure S2.**

Zoomed-in view of p97 N-terminal domains seen on the map with SPI bound to subunit B with the atomic models docked in. All six N-domains of the p97 hexamer are in “up” conformation.

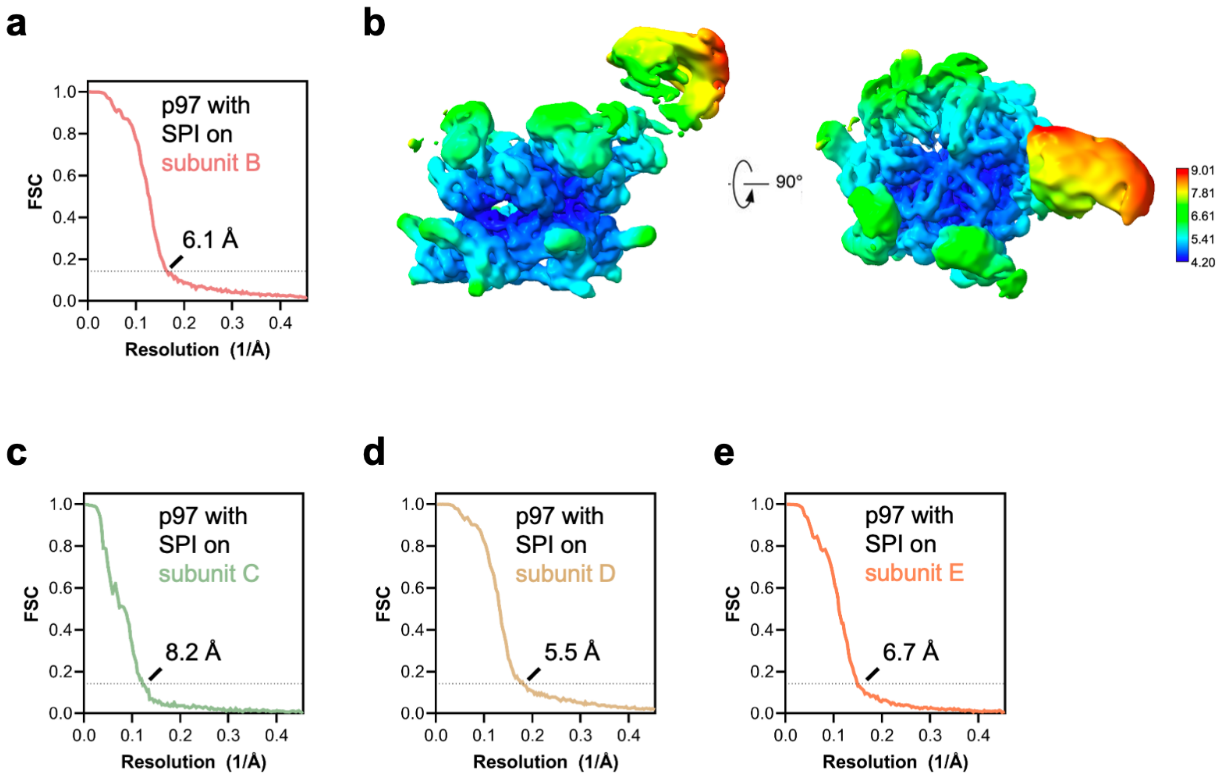


**Figure S3.**

(a) Gold-standard Fourier shell correlation (FSC) plot of the p97 map with SPI bound on the B subunit. For this map 191477 particles were used. (b) Local resolution of the p97 map with SPI bound on the B colored according to the scale on the right. (c) Gold-standard FSC plot of the p97 map with SPI bound on the C subunit. For this map 23526 particles were used. (d) Gold-standard FSC plot of the p97 map with SPI bound on the D subunit. For this map 253882 particles were used. (e) Gold standard FSC plot of the p97 map with SPI bound on the E subunit. For this map 82968 particles were used. Resolutions were determined at FSC = 0.143 (dotted line).

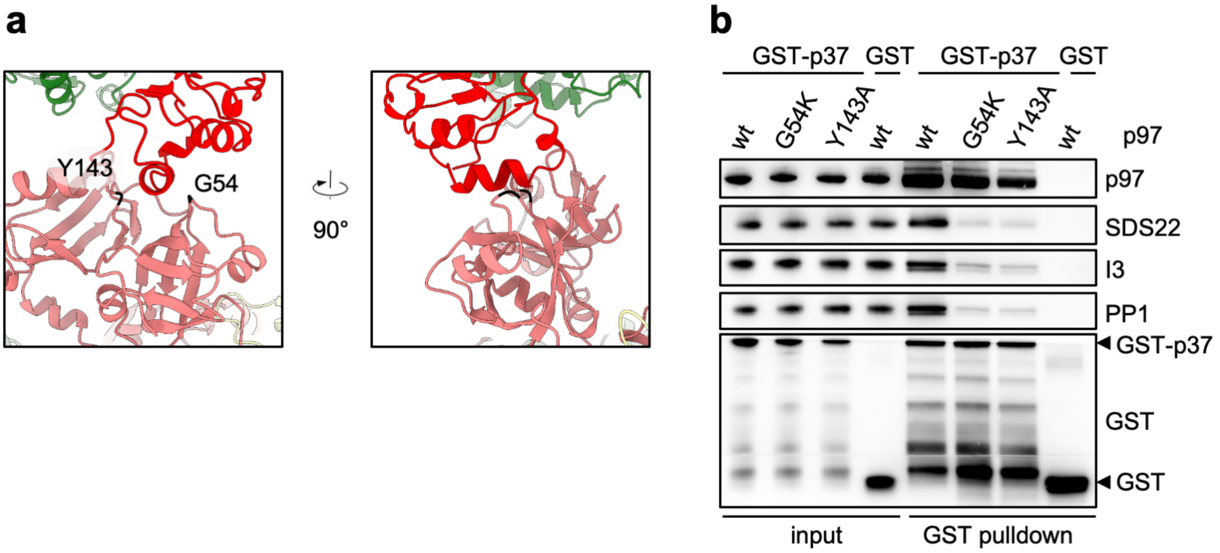


**Figure S4.**

(a) Views of the interaction site between SDS22 and the p97 N-domain groove. Mutated residues chosen for proximity to the SDS22-p97 contact site are labelled. (b) GST-pulldown binding assays using purified GST or GST-p37, SPI and p97 variants with the chosen mutations as indicated were carried out in the presence of ATP. Note that the mutations largely reduce SPI but not p37 binding.
